# Cell-free screening, production and animal testing of a STI-related chlamydial major outer membrane protein supported in Nanolipoproteins

**DOI:** 10.1101/2024.08.20.608363

**Authors:** Mariam Mohagheghi, Abisola Abisoye-Ogunniyan, Angela C. Evans, Alex Peterson, Greg Bude, Steven Hoang-Phou, Byron Dillon Vannest, Dominique Hall, Amy Rasley, Nicholas Fischer, Wei He, Beverly Robinson, Sukumar Pal, Anatoli Slepenkin, Luis de la Maza, Matthew A. Coleman

**Author notes:** Suggested as shared first co-authors (Mariam Mohagheghi, Abisola Abisoye-Ogunniyan).

## Abstract

Vaccine development against Chlamydia, a prevalent sexually transmitted infection (STI), is imperative due to its global public health impact. However, significant challenges arise in the production of effective subunit vaccine based on recombinant protein antigens, particularly with membrane proteins like the Major Outer Membrane Protein (MOMP). Cell-free protein synthesis (CFPS) technology is an attractive method to address these challenges as a method of high-throughput membrane protein and protein complex production coupled with nanolipoprotein particles (NLPs). NLPs provides a supporting scaffold while allowing easy adjuvant addition during formulation. Over the last decade we have been working toward production and characterization of MOMP-NLP complexes for vaccine testing. The work presented here highlights the expression and biophysical analyses, including transmission electron microscopy (TEM) and dynamic light scattering (DLS), confirm formation and functionality of MOMP-NLP complexes for use in animal studies. Moreover, immunization studies in preclinical models compare the past and present protective efficacy of MOMP-NLP formulations, particularly when co-adjuvanted with CpG and FSL1. Ex vivo assessments further highlight the immunomodulatory effects of MOMP-NLP vaccinations, emphasizing their potential in eliciting robust immune responses. However, further research is warranted to further optimize vaccine formulations, validate efficacy against *Chlamydia trachomatis*, and better understand underlying mechanisms of immune response.

## Introduction

*Chlamydia trachomatis* is an increasing global public health concern, affecting millions^1^ as one of the most common STIs, and is responsible for numerous clinical manifestations^2,7^. Chlamydia infection is often asymptomatic and can result in serious lifelong health complications, including pelvic inflammatory disease (PID), infertility, and ectopic pregnancy. Certain strains can also cause trachoma, leading to vision impairment^3^.

There are 18 different serovars of Chlamydia trachomatis, associated with three distinct clinical manifestations: (genital infections, ocular infections such as trachoma and lymphogranuloma venereum, and respiratory infections primarily caused by *Chlamydia pneumoniae*^6^. Serovars A-C results in serious ocular impairment by chronic conjunctivitis; Serovars D-K cause genital tract and neonatal infections^4,5^; Serovars L1-L3 manifest as an ulcerative disease of the genital area associated with genital ulcers or papules that may vary in severity^6^. Timely and effective treatment is crucial not only to alleviate symptoms but prevent significant complications and further infection. Current treatments include antibiotics, which are generally curative but have several limitations, such as the potential for antibiotic resistance and re-infection. Importantly, antibiotics do not provide cross-protection against other STIs, such as gonorrhea or syphilis^2,7^. Developing an effective vaccine could reduce population reliance on antibiotics and slow emergence of antibiotic resistance^2,8^.

*Chlamydia trachomatis* belongs to the *Chlamydiaceae* family and is a gram-negative, anaerobic, obligate intracellular organism that replicates within eukaryotic cells^9^. There is a distinct biphasic lifecycle characterized by two morphological forms: metabolically inactive extracellular elementary bodies (EBs) and metabolically active intracellular reticulate bodies (RBs)^4,10^. Membrane proteins play a pivotal role in mediating the interactions between Chlamydia and host cells, facilitating EB entry into host cells and the subsequent differentiation into metabolically active RBs^11^. Furthermore, membrane proteins participate in host cell manipulation, nutrient acquisition, and evasion of host immune responses^9^. Understanding these dynamics provides insight into Chlamydia pathogenesis and informs the development of effective intervention strategies. *Chlamydia muridarum*, the mouse-specific analog to human *Chlamydia trachomatis*, is used to study immune responses.I Its pathogenicity in mice and ability to model human female genital tract infections make it particularly useful for such studies.^12^ In particular, previous studies have shown that the major outer membrane protein (MOMP) of Chlamydia has been identified as a promising vaccine candidate^13,14^. MOMP, highly conserved across Chlamydia species, forms a trimeric structure that is membrane-bound. It accounts for over 60% of the rigid, outer-membrane surface of EBs, playing a crucial role in the pathogen’s structure and immune interactions (REF-Cardwell 1981). MOMP is highly immunogenic, containing numerous B- and T-cell epitopes that are critical for immune recognition. However, its production is challenging due to structural aggregation and insolubility^15-17^.

Cell-free protein synthesis (CFPS) allows for transcription and co-translation of otherwise insoluble membrane proteins, including chlamydial membrane proteins such as MOMP, polymorphic membrane, inclusion body proteins, translocase III proteins within nanolipoprotein particles (NLPs)^18,19^. The traditional NLP assembly method requires purified scaffold proteins, detergent-solubilized membrane proteins, and lipids, providing precise control over NLP composition. This control makes NLPs an ideal platform for targeted and tailored vaccine design^20-22^. We use a faster approach that combines apolipoprotein-encoded and membrane-protein-encoded DNA with lipids in a cell-free reaction chamber incubated at 25-30°C for 12-18 hours. This approach eliminates the need for detergents or pre-purified proteins. Additionally, empty NLPs can be produced, purified, and lyophilized to support future protein production and purification processes^23^.

CFPS-produced and purified MOMP-incorporated NLPs are a promising vaccine candidate. Previous studies have tested the protective properties of MOMP-NLP purified with a cell-free expression system^21^. MOMP-NLP vaccine formulations elicited robust humoral and cellular immune responses in preclinical mouse models, demonstrating significant induction of MOMP-specific antibodies and cytotoxic T lymphocyte (CTL) responses against Chlamydia infection^22^. Various studies have also explored recombinant MOMP vaccines with distinct delivery systems and adjuvants. For instance, a study by *Sahu et al*. demonstrated that PLA-PEG nanoparticles encapsulating recombinant MOMP induced robust antigen-specific Th1 and Th17 cytokine responses, alongside the proliferation of CD4+ T-cells and the development of effector T-cell phenotypes. This resulted in protective immunity against genital *Chlamydia muridarum* challenges in mice^24^. Additionally, in a study by *Tu et al*., showed that a MOMP-based vaccine elicited strong humoral and cellular immune responses to *C. trachomatis*, generating specific antibodies and cytotoxic T lymphocytes in a mouse model^25^.

Using NLPs as carriers for vaccines, particularly through our research that summarizes past and present formulations based on MOMP-NLPs, offers insight that help highlight formulations for the developing and testing Chlamydia vaccines. Consolidation of current data shows the advantages of NLP formulations include rapid production in cell-free expression, the capacity to incorporate additional components such as adjuvants, and the ability to mimic the native environment of a membrane-bound protein^23^. By incorporating adjuvants and other immunomodulatory agents, our approach with MOMP-NLPs can enhance immune efficacy against Chlamydia^21^. There is substantial potential of NLPs as a platform for presenting chlamydial antigens, including MOMP, offering a promising avenue to advance vaccine development. Here, we summarize both new and previously published data on using of cell-free NLP techniques for producting and testing MOMP-NLP formulations. Importantly, we consolidate and analyze MOMP-related vaccine data from in vivo and ex vivo experiments.

## Results

### 1. Nanolipoproteins (NLPs) are a tool for studying membrane bound proteins

NLP assemblies can be created using cell-free methods with or without membrane proteins to support both in vitro and in vivo studies (Figure 1). An NLP is an 8-25nm membrane-mimetic disc composed of phospholipids and a stabilizing scaffold protein, such as apolipoproteins (Figure 1a). Combining the apolipoprotein-encoded plasmid and lipid into a cell-free reaction yields “empty” NLPs without an associated membrane protein (Figure 1b). Cargo, such as a desired membrane protein, can be efficiently incorporated using lipids and plasmids that encode both the scaffold and membrane proteins in the same cell-free reaction. This reaction results in the incorporation of the full-length membrane protein into the assembled NLP (Figure 1c). Yields range from micrograms to milligrams of membrane protein-containing NLPs, depending on factors such as the scale of reaction, plasmid backbones for protein components, and the types of associated membrane proteins.

**Figure 1.**
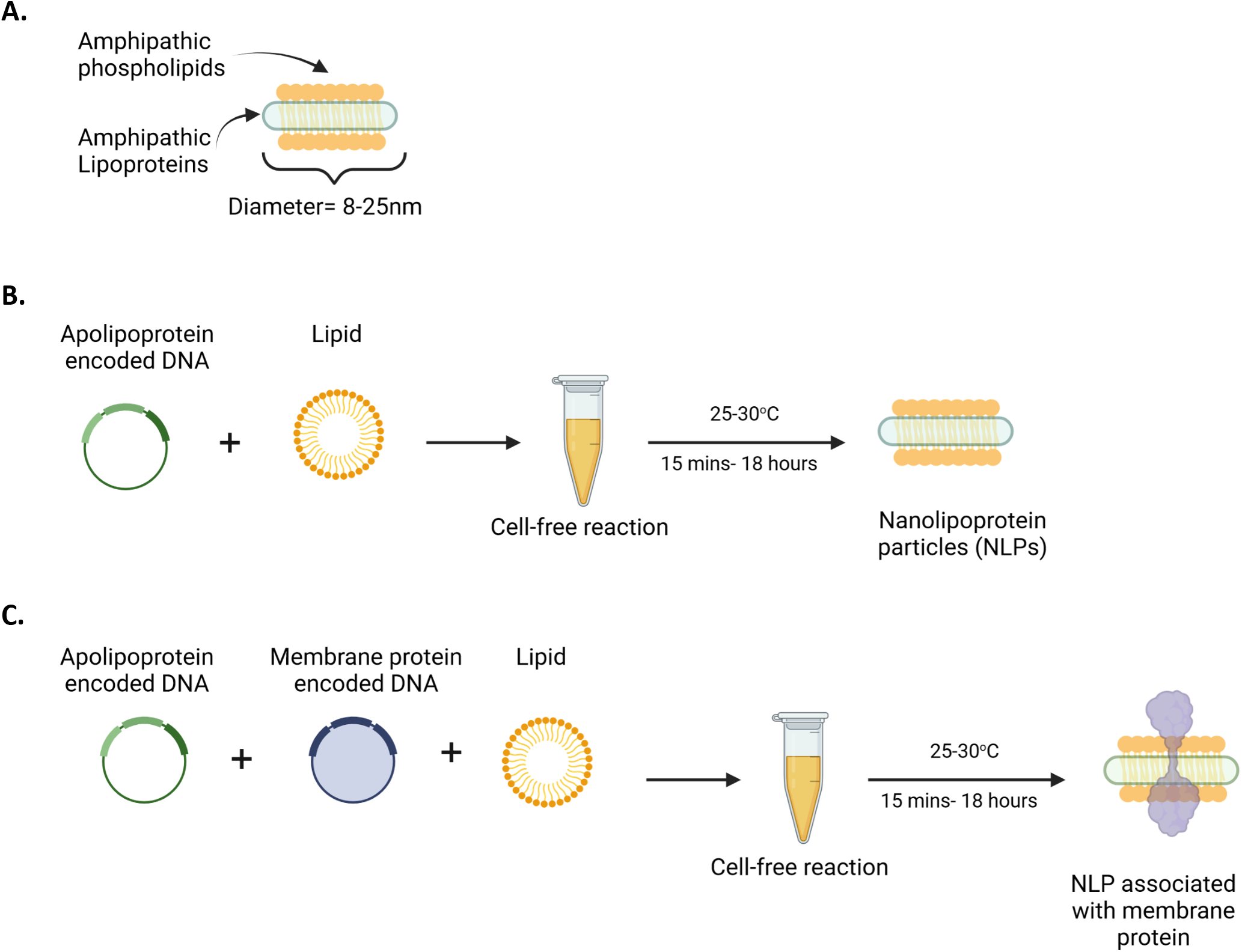
Nanolipoproteins are a tool for studying membrane bound proteins. (A) Nanolipoprotein particles, NLPs (8 - 25nm disc-shaped particles) formed by the spontaneous assembly of phospholipids into a bilayer stabilized by an apolipoprotein scaffold protein. (B) Cell-free approach for NLP production without detergents or pre-purified proteins. (C) Cell-free approach for full length membrane proteins expression encapsulated in NLPs.

### 2. Cell-free screening enables rapid lipid testing and optimization

To optimize the solubility of MOMP in NLPs, we screened 12 different lipid/polymer NLP formulations. Each condition was run as a 25 µL, *E. coli*-based cell-free reaction with the scaffold protein ApoA1. Specifically, we screened EggPC, DOPE, DOPC, and DMPC lipids for soluble MOMP after co-translation with ApoA1 and incorporation of fluorescently labeled amino acids (Figure 2a). We also combined these lipids with telodendrimer PEG^5000^-CA_8_ (Telo) in our screen, which has been shown to increase complex solubility^22^. Out of the four individually tested lipids, DMPC performed the best, indicated by denser bands at 40kDa on the gel, representing more MOMP-encapsulated NLP compared to those with EggPC, DOPE, and DOPC. Notably, the addition of telodendrimer PEG^5000^-CA_8_ (Telo) to DMPC reactions not only enhanced the total MOMP-NLP complex but also improved solubility. Furthermore, other conditions of the cell-free reaction such as reaction volume, membrane/scaffold protein plasmids, reaction time, and temperature can be systematically investigated to optimize the production of soluble membrane-protein-containing NLP. For example, in a 100 µL *E. coli*-based cell-free reaction, we demonstrated optimization of the NLP scaffold protein (Figure 2b). Addition of ApoA1 scaffold protein-plasmid results in stronger MOMP-NLP bands in comparison to ApoE4 scaffold protein plasmid. However, omitting a scaffold protein entirely dramatically reduces MOMP-NLP solubility to nearly undetectable levels compared to ApoA1 and ApoE4 scaffold proteins. Combining these findings with previous lipid-screen results, we identified the most effective formulation for soluble MOMP-NLP complex is a combination of ApoA1 scaffold protein with DMPC and Telo. This formulation is ideal for scale-up, purification and characterization in support of in vivo vaccine-related studies.

**Figure 2.**
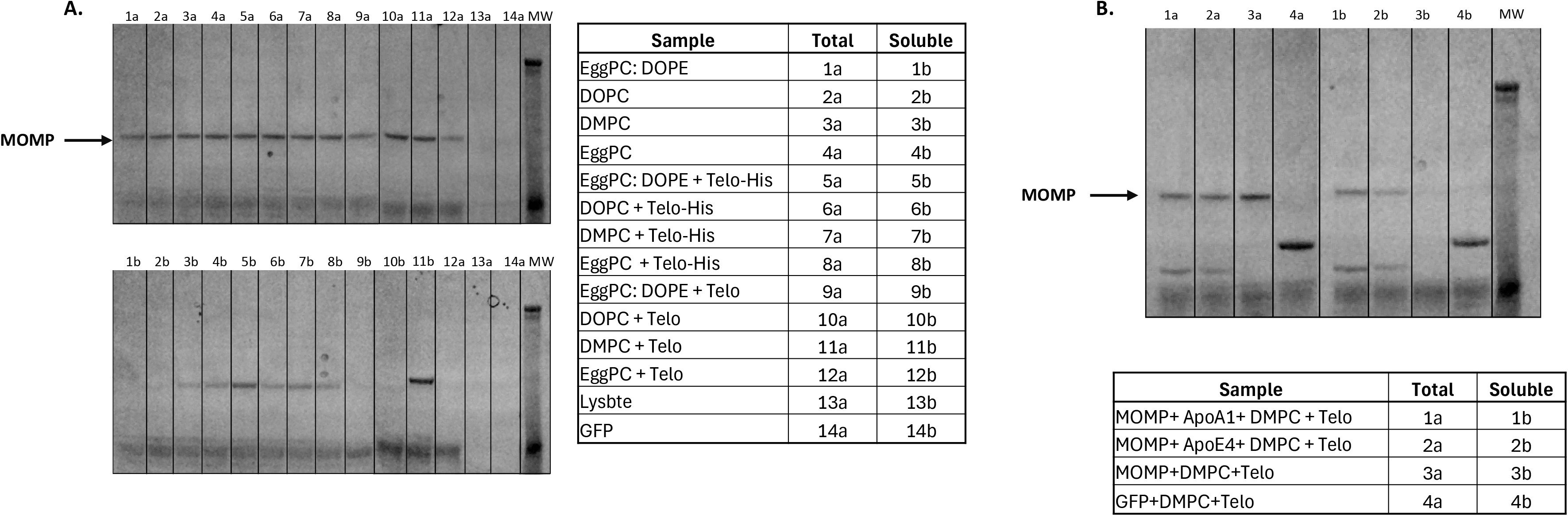
Cell-free screening allows for rapid, large-scale condition testing and optimization. (A) SDS –PAGE image of 25 µL MOMP protein reactions with various lipids and telodendrimer to identify condition that optimizes solubility. (B) SDS page of 100 µL MOMP protein reactions with apolipoprotein scaffold, and various lipid/telodendrimer combinations demonstrates scalability of the cell-free techniques. M = Molecular weight marker.

### 3. Cell-free production and purification of MOMP-NLP protein complex

We first reported MOMP solubilization and characterization in NLPS using cell-free techniques in the Wei et al publication in 2016. In addition to serving as a viable method for rapid formulation screening, cell-free reactions are scalable to produce sufficient yields of MOMP-NLP for in vivo immune studies^26^. For cell-free production of MOMP-NLP, we increased the *E*.*coli*-based reaction volume to 1-2 mL. A detailed protocol for cell-free MOMP-NLP production can be found in the Materials and Methods section. Briefly, self-assembly of MOMP-NLP complexes is carried out in a cell-free reaction by combining MOMP and ApoA1 protein-encoded plasmids with lipids and telodendrimers in a *E*.*coli*-based cell lysate reaction mix (Figure 3a). Affinity purification with Ni^2+-^NTA-resin results in two distinct bands on the SDS-PAGE gel, confirming two proteins in the complex, MOMP and ApoA1. Unlike the ApoA1 protein, the translated MOMP protein lacks a His-tag, which indicates that MOMP is incorporated into the NLP and co-purified with it as part of the membrane-protein NLP complex. Quantification and densitometry of affinity purified and dialyzed MOMP-NLPs was performed, indicating MOMP protein to ApoA1 scaffold protein ratios range from 1:1 to 3:1. A typical 1 mL reaction yields 1.5-2 mgs of total protein per mL, including both MOMP and ApoA1 proteins (Figure 3b). Following dialysis in Tris-based buffers, yields range from 300-500 µg of pure MOMP membrane protein in the MOMP-NLP complex for downstream use. Thus representing 30 to 50 potential doses of vaccine per mL of reaction volume. Characterization of purified and dialyzed MOMP-NLP complexes by Western blot with the MOMP-specific mAb40 antibody (Figure 3c), which targets a linear epitope located on the extracellular domain of the MOMP protein^21^ confirms the protein co-purified with NLPs is MOMP. Taken together, these results indicate that cell-free production of MOMP-NLPs is readily purifiable via affinity chromatography and may be subsequently used in downstream biophysical characterization, in vitro, and in vivo assays.

**Figure 3.**
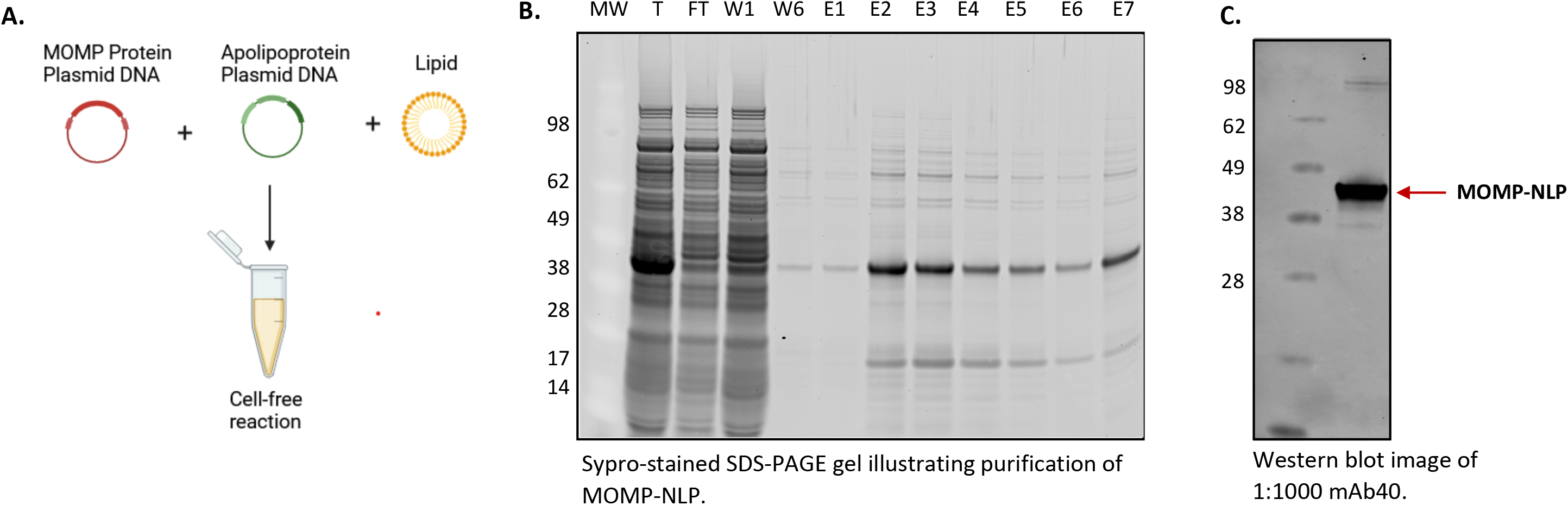
Cell-free production of MOMP-NLP protein complex for mice studies. (A) Cell-free expression of MOMP-NLP showing components including MOMP DNA, Δ49ApoA1 DNA and pre-prepared DMPC/telodendrimer lipid in a cell-free reaction chamber. (B) SDS-PAGE Sypro-stained image showing cell-free produced MOMP-NLP purified using Nickel bead gravity column (molecular weight (MW) marker-SeeBlue2Plus, Total protein (T), flow through (FT)-MOMP protein not associated with an NLP, two of six washes (W)- to purify protein of interest, and seven elutions (E1-E7)- to recover MOMP-NLP. (C) Western blot micrograph showing cell-free produced MOMP using mAb40, a primary antibody against MOMP.

### 4. Characterization of the MOMP-NLP complex

Biophysical methods are used to ensure the formation of functional MOMP-NLP oligomeric complex. Negative stain cryo-TEM image analysis of the peak MOMP-NLP elution fractions (Figure 4a) confirms the association and insertion of MOMP protein into the NLP. Multiple MOMP proteins can be incorporated in one nanodisc, representative of the native trimer formation of MOMP. Additionally, dynamic light scattering (DLS) supports insertion of MOMP protein into NLPs. Empty NLPs (not shown) measure at approximately 10 nanometers, whereas MOMP-NLPs have a size closer to 45nm, plus the standard deviation (Figure 4b)^22^. Electrophysiology assays were used to confirm that the MOMP-NLP protein forms a functionally active porin within lipid bilayers (Figure 4c). The electrophysiology current trace indicates an open pore (top), shown by a 10pA jump. A flat trace (bottom) suggests a lack of pore function, either due to the absence of the inserted porin or a closed pore (Figure 4d). Additional characterization by size-exclusion chromatography (SEC) confirmed homogeneity of MOMP-NLP complexes, as indicated by uniform peak elution fractions (data not shown). In previous experiments, the MOMP-NLP complex eluted at approximately 7 minutes, indicating that the complex can be separated from free proteins or lipid aggregates^22^.

**Figure 4.**
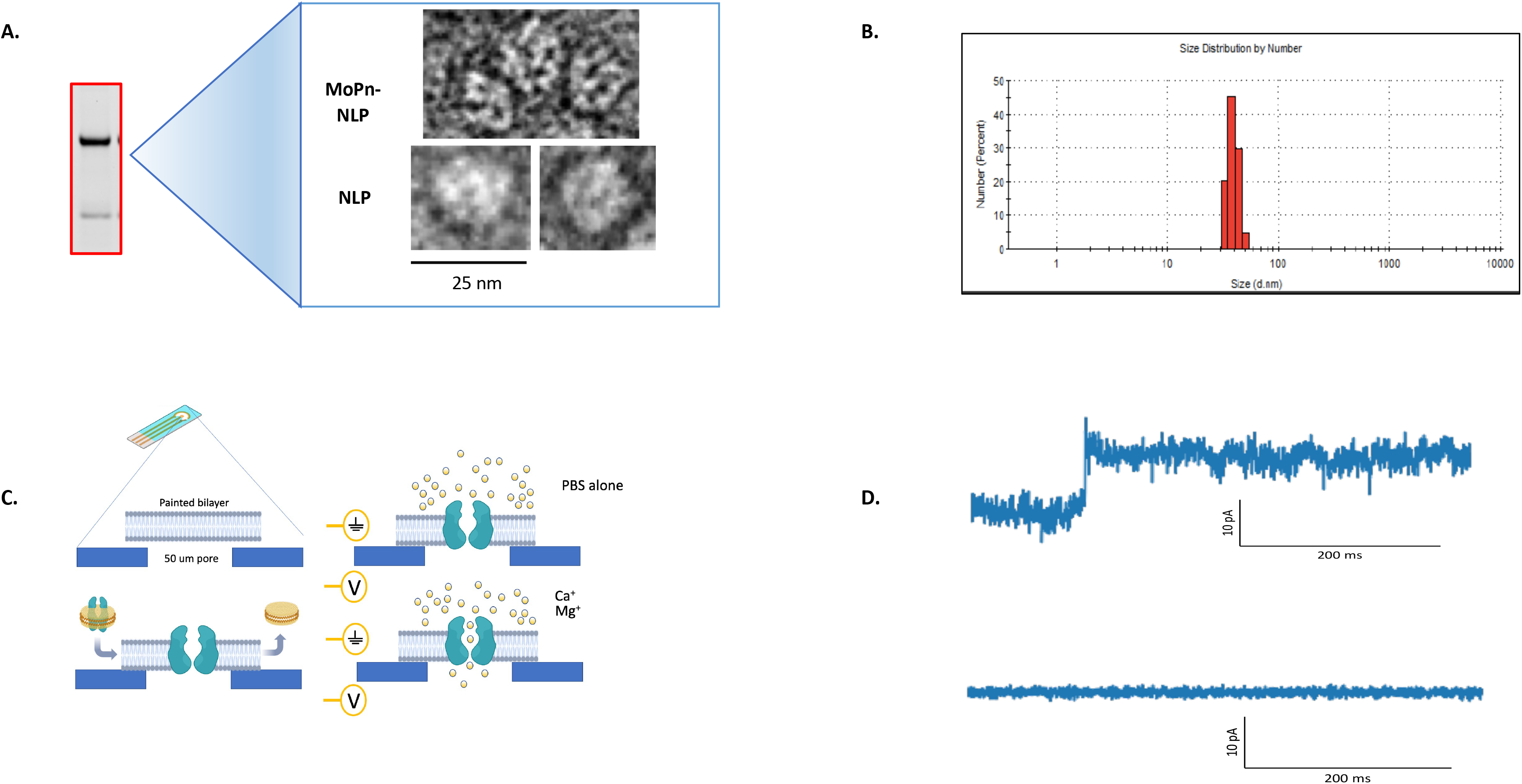
Imaging and electrophysiology techniques confirm formation of MOMP-NLP complex. (A) Negative stain cryoEM of peak fraction of eluted, soluble MOMP-NLP complex (from figure 3B) illustrates the circular disk-like shape of *C. muridarum* MOMP-NLP and empty NLP. (B) Dynamic Light Scattering (DLS) measurement further confirms formation of MOMP-NLP complex. (C) The MOMP protein is active as a porin as shown by electrophysiology. Single channel conductance assay of MOMP-NLPs in fixed bilayers at fixed voltage, standard 200mV. (D) Electrophysiology additionally confirms MOMP-porin is active; top trace shows 10 picoamp jump confirming functional porin activity. Flat line indicates no porin activity and baseline membrane bilayer reading value.

### 5. Effectiveness of NLP encapsulated MOMP-based vaccine formulations administered through various routes and immunization schedules

To test the effectiveness of our NLP encapsulated MOMP-based vaccine formulations, we explored several routes of vaccine administration-intranasal (IN) and intramuscular (IM)- along with both prime and prime-boost vaccination regimens in Balb/c female mice (Figures 5 and 6). To complement these studies, we also explored a prime-boost-boost vaccination schedule and used ex vivo splenocyte cultures to assess the immune-relevant pathways (Figure 7).

**Figure 5.**
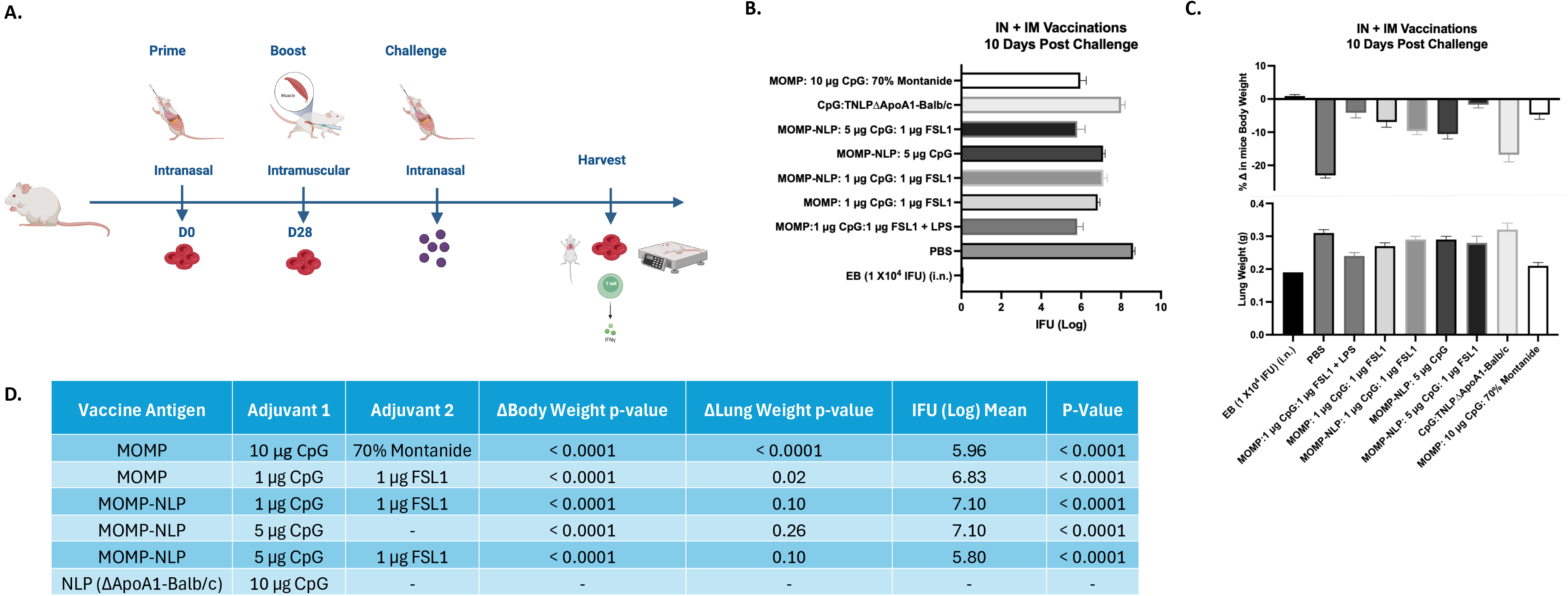
Systemic and local disease burden following intranasal *C. muridarum* challenge of mice vaccinated via IN/IM prime-boost regimen with recombinant MOMP-NLP, and adjuvanted with CpG and FSL1. (A) Experimental schematic showing an intranasal prime **and** intramuscular boost vaccinations, i.n. challenge and tissue harvest. (B) Log IFU of *C. muridarum* recovered from mice lungs 10 days post challenge. (C) Change in mice body weight and lung weight (g) at 10 days post i.n. challenge with *C. muridarum*. (D) Summary chart showing vaccine antigen, adjuvants, and *p values*.

**Figure 6.**
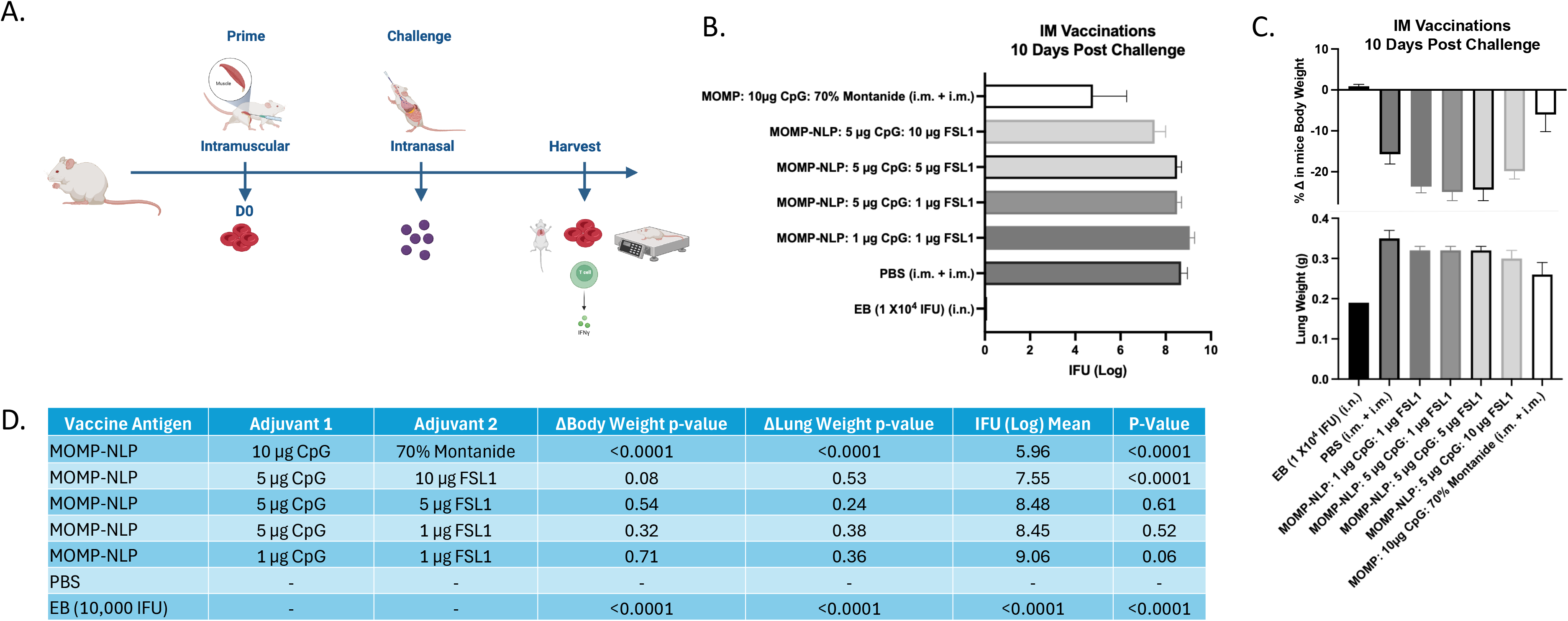
Systemic and local disease burden following intranasal *C. muridarum* challenge of mice vaccinated via IM prime regimen with recombinant MOMP-NLP, and adjuvanted with CpG and FSL1. (A) Experimental schematic showing intramuscular prime vaccinations, i.n. challenge and tissue harvest. (B) Log IFU of *C. muridarum* recovered from mice lungs 10 days post challenge. (C) Change in mice body weight and lung weight (g) at 10 days post i.n. challenge with *C. muridarum*. (D) Summary chart showing vaccine antigen, adjuvants, and *p values*.

**Figure 7.**
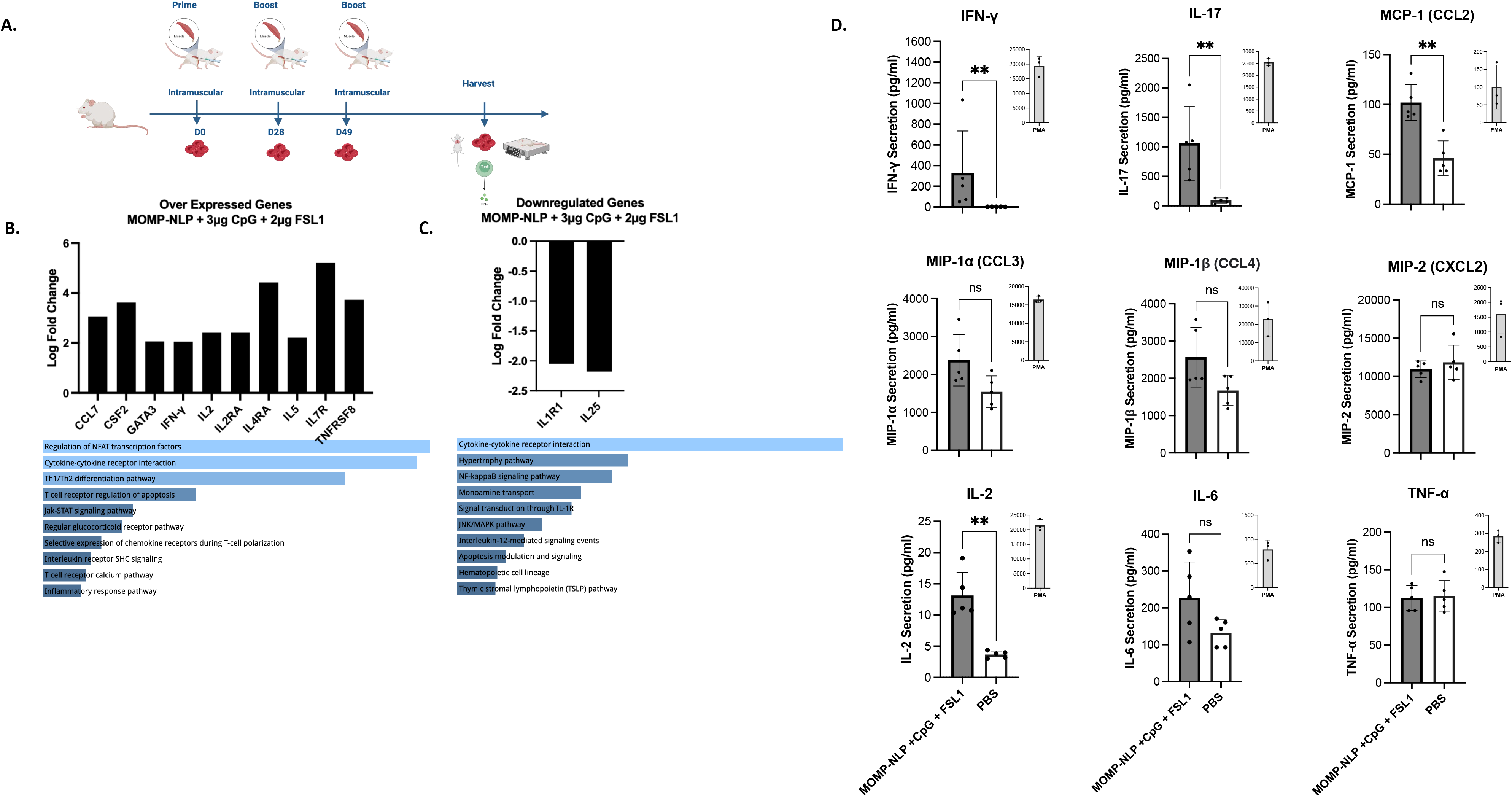
Gene and cytokine expression following prime-boost-boost mice vaccinations with recombinant MOMP-NLP adjuvanted with CpG and FSL1. (A) Experimental schematic showing an intramuscular prime and two intramuscular boost vaccinations followed by tissue harvest. (B) Overexpressed genes with greater than or equal to 2-fold change value of vaccinated mice relative to sham (PBS) vaccinated mice and respective activated pathways, ranked by p–value. (C) Illustrates the downregulated genes. For these experiments, cDNA samples were obtained from pooled spleen RNA samples of 5 different animals per group. (D) Secreted cytokines following stimulation of splenocytes 7-10 days after boost with MOMP. *(P value ranked pathways / processes generated by NCATS BioPlanet, Enrichr)*

In vivo studies relied on intranasal challenge studies with C. *muridarum* four weeks after the last vaccination, followed by tissue harvest 10 days post-challenge (Figures 5a and 6a). Systemic and local disease burdens following an intranasal *C. muridarum* challenge in all vaccinated mice were evaluated by measuring body weight, lung weight and the log IFU of *C. muridarum* recovered from the lungs for all the vaccination schedules. Mice vaccinated using the IN/IM prime-boost regimen with recombinant MOMP-NLP adjuvanted with 5 µg CpG and 1 µg FSL1 showed more than a 1-log fold reduction of *C. muridarum* IFUs recovered from the lungs (mean of 5.8 log IFU, mean) compared to mice vaccinated with the same schedule and a lower dose of CpG, 1 µg (mean of 7.1 log IFU), and almost a 3 log fold reduction compared to those recovered from mice vaccinated with a sham control of PBS (mean of 8.6 log IFU) (Figure 5b). Interestingly, our single adjuvanted recombinant MOMP-NLP formulation with only 5 µg of CpG offered the same level of protection (mean of 7.1 log IFU) as the formulation combining 1 µg of CpG with 1 µg of FSL (mean of 7.1 log IFU) (Figure 5b). Additionally, mice vaccinated with MOMP-NLP adjuvanted with 5 µg CpG and 1 µg FSL1 showed a significantly lower change in body weight and reduced lung weight when compared to the PBS vaccinated mice (Figure 5c). All mice vaccinated with either a MOMP or MOMP-NLP based formulation showed a significant reduction of *C. muridarum* IFUs recovered from their lungs compared to mice vaccinated with PBS (Figure 5d).

Although a combined IN/IM vaccination scheme shows promising results for a protective vaccine, we also wanted to understand the effectiveness of our MOMP-NLP antigens administered solely through the IM vaccination route. For this vaccination scheme, we tested our MOMP-NLPs along with varied concentrations of CpG and FSL1 adjuvants in Balb/c female mice with, an IM prime vaccination followed by an intranasal challenge with *C. muridarum* (Figure 6a). The MOMP-NLP antigen adjuvanted with 5 µg CpG and 10 µg FSL1 showed the highest level of protection, with a mean of 7.5 log IFU of *C. muridarum* recovered from the lungs of mice compared to a mean of 9.06 log IFU for the formulation with lower doses of the adjuvants (1 µg CpG and 1 µg FSL1) (Figure 6b). Although not significant, the change in body weight and lung weight were lower for the formulation with 5 µg CpG and 10 µg FSL1compared to those with other tested concentrations of CpG and FSL1(Figure 6c and Table 6d).

Lastly, we aimed to characterize how MOMP-NLP antigens and various adjuvant combinations interact with and induce the immune system to potentially elucidate the currently unknown correlates of protection against Chlamydia infections. To accomplish this, we injected MOMP-NLPs adjuvanted with 3µg CpG and 2µg FSL1 into Balb/c female mice following a prime-boost-boost vaccination regimen without intranasal challenges (Figure 7a). Gene expression of pooled spleen samples of MOMP-NLP + CpG + FSL1 vaccinated mice, compared to those from PBS vaccinated mice, showed overexpression of Th1 genes-IFNγ, IL2, IL2RA, IL4RA, and Th2 genes-IL5, CCL7, CSF2, GATA3, IL7R and TNFRSF8. These genes are involved in the activation of several pathways including the regulation of the transcription factor-nuclear factor of activated T cells (NFAT), cytokine-cytokine receptor interaction and Th1/Th2 differentiation genes (Figure 7b). Concurrently, the genes IL1R1 and IL25, which are involved in the mediating cytokine-induced immune and inflammatory responses and in inducing of NF-kappaB activation, respectively were downregulated (Figure 7c). Th1 and Th17 cytokines, IFNγ, CCL2 and IL17 were significantly secreted in the conditioned media of splenocytes from MOMP-NLP + CpG + FSL1 vaccinated mice cultured *ex vivo* and restimulated with recombinant MOMP compared with PBS restimulated controls (Figure 7d).

## Discussion

Combining the NLP technology with an *E. Coli* cell-free expression system aids rapid membrane protein synthesis, maintaining the native oligomeric formation necessary for effective immunogenicity^27^, and serves as a powerful tool for developing vaccines based on membrane-bound proteins representing antigens derived from of infectious agents. NLPs combined with cell-free expression systems^19,28,29^, provide a rapid and flexible approach to produce a wide range of difficult-to-produce proteins in a format that allows for quick screening of multiple conditions to optimize antigen solubility^30^. Another attractive feature of this technology is its ability to couple the process with vaccine studies, capitalizing on the NLP’s role as as a delivery vehicle.

Within this paper, we present a more complete view of the production of the MOMP-NLP complex^22^ using various adjuvant formulations for in vivo and ex vivo studies. Previous in vivo studies have demonstrated that the native MOMP structure is crucial to elicit a systemic immune response^21,22^. CombiningMOMP and apolipoprotein plasmid-DNA with a lipid and Telo rapidly produces the MOMP-NLP complex. Previous TEM imaging studies have shown that the addition of the Telo plus lipids not only l serves as support for membrane proteins forming a unique type of nanodisc^22^, but may aid in solubilizing lipids, thereby facilitating the formation of MOMP-NLP^22^. It is also hypothesized that the PEGylated tail minimizes MOMP and MOMP-NLP interactions, thus reducing protein aggregation and retaining water solubility. The MOMP-NLP complex and its trimeric formation is confirmed through several quality control checks and structural analyses, including size exclusion chromatography (SEC), TEM, SDS-PAGE, and dynamic light scattering^21,22,26^. Previous reports indicate that native trimeric MOMP is the optimal antigen to induce a protective immune response^31,32^. Electrophysiology results shows that the expression of MOMP in NLPs forms a complex with MOMP, potentially with a percentage of the protein presenting as a functional trimeric channel, thus allowing it to oligomerize in a manner similar to native MOMP^33^. However, these previous studies did not use native MOMP as a control in electrophysiology experiments.

Protective immunity involving both cellular and humoral immune responses following vaccination is the goal for vaccine development for infectious diseases^34^. Animal studies have demonstrated that recombinant MOMP can elicit long lasting and protective antibody responses specific to *C. muridarum*^*35*^. Hence, the expression of MOMP in NLPs presents a promising vaccine delivery platform that has the potential to enhance targeted antigen presentation and the colocalization of different adjuvants with MOMP for robust immunogenicity^36^. We previously demonstrated that mice vaccinated via the IN/IM prime-boost regimen with MOMP-NLP and CpG and FSL1 as adjuvants elicited partial protection after intranasal challenge with *C. muridarum*^*21*^. In this study, we tested different concentrations of CpG in formulation with recombinant MOMP-NLP and FSL1 and explored different routes of vaccination. Our formulation with the higher dose of CpG administered using the IN/IM prime-boost regimen showed significantly fewer *C. muridarum* IFUs in the lungs of mice, with minimal change in body weight, indicating a potent formulation resulting in the improvement of the quality and quantity of immune responses^37,38^ to our recombinant MOMP-NLP formulation, generated by using CpG and FSL1 as co-adjuvants.

To understand the immunogenicity of our *E. Coli* cell-free expressed MOMP-NLP complex in combination with CpG and FSL1, we utilized unchallenged, IM prime-boost-boost vaccinated mice and measured the gene expression profiles and *ex vivo* cytokine and chemokine secretion from spleen RNA and splenocytes, respectively. These IM vaccinations were three weeks apart and tissue harvest was carried out 7 - 10 days after the last boost. The genes overexpressed by the MOMP-NLP adjuvanted with CpG and FSL1 significantly regulate NFAT (Nuclear Factor of Activated T-cells) transcription factors, which are known for playing a crucial role in the regulation of genes involved in T-cell activation, proliferation, and differentiation. They are activated upon T-cell receptor stimulation, ultimately influencing immune responses including inflammation, immunity, and tolerance^39,40^. Cytokine-cytokine receptor interactions were also activated. These interactions play a crucial role in the inducing immune responses by mediating communication between immune cells and other tissues^41,42^. Furthermore, we observed activation of the Th1/Th2 differentiation pathway, where naïve CD4+ T helper cells are polarized to either Th1 or Th2 cells, orchestrating appropriate immune responses against pathogens and antigens^43,44^. The two genes downregulated with MOMP-NLP adjuvanted with CpG and FSL1 vaccination, IL1R1 and IL25, are involved in cytokine-cytokine receptor interactions as well. ILIR1 is a mediator of many cytokine-induced immune and inflammatory responses, while IL25 has been shown to be associated with type 2 immune responses responsible for the development of allergic diseases^45,46^.

*E. Coli* cell-free expression is a powerful technique that allows for rapid screening and optimization of difficult-to-produce Chlamydial membrane proteins. The technology can be coupled with the NLP platform to increase membrane protein solubility and yield. This allows for the formation of the functional, multimeric MOMP-NLP complex whose structure and formulation characteristics can be analyzed using biochemical techniques. Our approach focuses on techniques such SDS-PAGE, Western blots, TEM, DLS, and electrophysiology. The NLP-based approach provides highly purified and stable material for characterization and utility in animal studies. Overall, we have been able to demonstrate that *C. muridarum* MOMP formulated in a biomimetic nanoparticle is malleable for the colocalization of different adjuvants that can enhance the robust, protective, and prolonged immune responses against *Chlamydia* infections.

While our past and present studies have demonstrated the efficacy and immunogenicity of our MOMP-NLP based formulations in preclinical models of Chlamydia infection, there are several aspects that remain unexplored. Firstly, there is a need to directly compare the folding and functional properties of our MOMP-NLP complex to isolated native MOMP protein. Although our characterization assays (SDS-PAGE, Western blotting, TEM imaging and electrophysiology) suggest the formation of a functional oligomeric complex, further structure analyses, such as X-ray crystallography or cryo-electron microscopy, are necessary to confirm the folded state of MOMP within the NLP. Additional investigation into the dose-response relationship of both MOMP and different adjuvants tested is crucial. Identifying the optimal dose of MOMP and adjuvants is key for maximizing vaccine efficacy while minimizing potential adverse effects. Furthermore, while our experiments have *Chlamydia muridarum* (Cm) MOMP as the antigen, translating these findings to *Chlamydia trachomatis* is essential. Despite conserved regions, additional studies are required to validate the immunogenicity and efficacy of our vaccine formulations against *C. trachomatis* infection. Overall, addressing these aspects will provide valuable insights into the development and optimization of MOMP-NLP based vaccines for Chlamydia infection.

## Conclusion

Presenting comprehensive data on the generation of NLP protein complexes is crucial. It allows for evaluating the potential and necessity of exploring additional chlamydial antigens and adjuvants that could enhance MOMP’s efficacy as a subunit vaccine. The consolidated findings from this research provide a unique dataset on the screening and production pipeline for MOMP-NLP vaccine development, potentially paving the way for other STI-related vaccines targeting membrane-bound antigens. The NLP platform serves as an optimal tool for the co-delivery of various lipidic adjuvant combinations. Notably, MOMP-NLP formulations with CpG and FSL1 demonstrated protective effects, identifying key immunological correlates essential for understanding protection mechanisms in mouse models. Continued research is imperative to optimize adjuvant dosages and combinations, aiming to improve protection against Chlamydia infections and reduce disease burden. A deeper understanding and strategic use of the mechanisms of action of these adjuvants, whether used individually or in combination, will be crucial in achieving the desired cellular and humoral immune responses necessary for developing a human Chlamydia vaccine.

## Materials and Methods

### Plasmids

Plasmids were previously described and published^21,22^. Briefly, codon-optimized sequences for the mouse ApoA1(Δ1-49), also known as Δ49ApoA1 gene and mouse MOMP (mMOMP)gene were assembled from oligonucleotides and cloned into Nde1/BamHI-digested pVEX2.4d vector (Roche Molecular Diagnostics) via Gibson Assembly as previously described^22^. The pIVEX24.d cloned with Δ49ApoA1 gene included a His tag used for downstream nickel affinity purification. The mMOMP gene-containing plasmid does not contain a His-tag.

### DMPC/telodendrimer preparation

Production and use of telodendrimer as a nanodelivery tool for membrane protein support has been described and previously published^47^. The general protocol is as follows: PEG^5000^-CA_8_ telodendrimer is stored in powder at 20°C and is reconstituted with UltraPure DI water to a final working concentration of 2mg/ml. Preparation of small unilamellar vesicles (SUVs) from 1,2-dimyristoyl-sn-glycero-3-phosphorylcholine (DMPC) lipid were sonicated using a qSonica Q500 and one-eighth inch probe tip. The DPMC (stored in powder at -20°C) was reconstituted with UltraPure DI H2O to a concentration of 20 mg/mL and sonicated using 15 seconds on and 15 seconds off pulse intervals at 22% amplitude for 2 minutes with a total input energy range of 600-700J. The sonicated lipids are centrifuged at 14,000 rcf for 1 minute to remove metal contamination from the probe tip. For the DMPC/ PEG^5000^-CA_8_ mixture, a 1:1 ratio of 20mg/mL DPMC and 2mg/mL PEG^5000^-CA_8_.

### Cell-free reaction

Scalable reactions (25 µL, 50 µL, and 1 mL) were performed with the RTS 500 Proteomaster *E. Coli* HY kit (Biotechrabbit GMbH, Hannover, Germany). Small-scale reactions contained the same ratio of components as large-scale reactions. Reaction components included lysate, reaction mixture, feeding mixture, amino acid mixture, and methionine and were added sequentially per manufacturer instructions. MOMP-NLP complex is produced by addition of 0.88 µg of delta49ApoA1 and 15 µg of mMOMP plasmid DNA to each 1 mL reaction. DMPC and telodendrimer are added in equivalent 100 µL volumes per reaction, for a total of 200 µL DMPC/telodendrimer mixture per 1 mL reaction. Reactions are incubated at 30°C shaking at 300 rpm for 16-18 hours.

### Affinity purification of NLP-related complexes

Gravity nickel affinity chromatography is used to isolate the MOMP-NLP complex from cell-free reaction mixture. 1 mL of a slurry of cOmplete His-Tag Purification Resin (Roche Molecular Diagnostics) was equilibrated with lysis buffer (50 mM NaH2PO4, 300 mM NaCl, pH 8.0) containing mM imidazole in a 10 mL chromatography column. The 1 mL cell-free reaction was mixed with equilibrated resin and incubated at 4°C for 1 hour. Post-incubation, the sample was washed with 1 mL of 20 mM imidazole buffer six times. MOMP-NLPs were eluted in six 300 µL fractions of buffer containing 250 mM imidazole and one final elution of 300 µL in 500 mM imidazole. Elutions were analyzed downstream via SDS-PAGE to identify peak fractions, which are further pooled and dialyzed into 50 mM Tris-HCl, 300mM NaCl, pH 7.5, and then stored at 4°C. Protein quantifications were performed using a Qubit instrumentation according to manufacturer’s instructions (ThermoFisher Scientific, Carlsbad CA). MOMP-NLP samples for mouse studies were tested for endotoxin levels with the Endosafe-PTS (Charles Rivery, Charleston, SC) endotoxin testing system based on Limulus amebocyte lysate. MOMP-NLP preparations had an average endotoxin between 1000-1500 endotoxin units per milliliter.

### Size exclusion chromatography (SEC)

NLPs are purified by SEC (Superdex 200, 3.2/300 GL column, GE Healthcare). SEC was run at a flow rate of 0.2 mL/min in PBS buffer with 0.25% PEG 2000.

### SDS-PAGE

1 - 5 µL aliquots of eluted MOMP-NLPS were mixed with 4x NuPAGE lithium dodecyl sulfate sample buffer and 10x NuPAGE sample reducing agent (Life Technologies), heat-denatured at 95°C for 5 minutes and loaded onto a 4-12% gradient premade 1.0 mM Bis-Tris gel (Life Technologies) with molecular weight standard SeeBlue Plus2 (Life Technologies). The running buffer was 1x MES-SDS (Life Technologies). Samples were run at 200V for 35 minutes. Gels were stained with SYPRO Ruby protein gel stain (Life Technologies) per manufacturer’s instructions and imaged using LI-COR Odyssey imager. Protein bands were quantified using Image Studio V2.0 software via densitometry.

### Western blot analyses

SDS-Page gel was run as previously described^22^. Briefly, the gel was transferred to PVDF membrane after running using the iBlot 2 Dry Blotting System (Invitrogen) for 10 minutes per manufacturer instructions. The membrane was then incubated at 4°C for 1 hour in LI-COR blocking buffer followed by an incubation in primary antibody (1:1,00 for mAb40) with LI-COR blocking buffer and 0.2% Tween 20 overnight at 4°C. The following day, the membrane was then washed for 5 minutes in 1x PBS-T (0.2% Tween 20, 1x PBS at pH 7.4), four times. The membrane was incubated and protected from light for 1 hour in secondary IR800 antibody (1:10,000 dilution) with blocking buffer and 0.2% Tween 20 at room temperature. Subsequently, 5-minute wash steps with 1x PBS-T were repeated four times. Afterwards, the membrane was washed with 1x PBS at room temperature to remove any residual detergent. Membranes were then imaged using LI-COR Odyssey imager.

### Dynamic light scattering (DLS)

Dynamic light scattering measurements of the NLP size were performed on a Zetasizer Nano ZS90 (Malvern Instruments, Malvern, UK) following the manufacturer’s protocols. Each data point represents an average of at least 10 individual runs.

### Animal vaccinations for immunological studies

Four to five-week-old female BALB/c (H-2d) mice (Charles River Laboratories; Wilmington, MA, USA) were housed at the Lawrence Livermore National Laboratory (LLNL) Vivarium. Animal protocols used were approved by the LLNL IACUC. The adjuvants CpG-1826 (TriLink, San Diego, CA, USA; 10g/mouse/immunization) and FSL-1 were directly mixed with single antigens (MOMP-NLP or MOMP at 10 μg of each antigen/mouse/ immunization) and different adjuvant combinations. Groups of 5 mice were immunized in a prime-boost-boost regimen, 3 weeks apart via the intramuscular (i.m.) route in the quadriceps muscle. To determine the cell-mediated immune responses all mice are euthanized 7-10 days post last boost and tissue of interest are harvested. All animal experiments were replicated at least once.

### RNA extraction and gene expression analysis

A fraction of the mouse spleen is collected at 7-10 days post last boost and stored in RNA*later* (Thermo Fisher). Total RNA is isolated using the column based nucleic acid purification kit (RNeasy, Qiagen), strictly according to the manufacturer’s protocol. Spleen samples were placed in reinforced polypropylene tubes containing ceramic beads (Omni Bead Ruptor tubes) and the appropriate lysis buffer. Samples were homogenized and subjected to a multi-step washing and centrifugation protocol. RNA purity and final concentration is determined followed by cDNA synthesis by reverse transcription (RT). PCR analysis was used to screen for changes in gene expression related to Th1 and Th2 responses. For this, the cDNA of 5 animals in each experimental group were pooled and mixed with the PCR master mix buffer (Rt^2^ Syber Green ROX qPCR Primer Assay, Qiagen) and analyzed on specific array plates (RT^2^ Profiler™ PCR Array Mouse Th1 & Th2 Responses-GeneGlobe ID - PAMM-034Z). PCR-array analysis was performed using a ViiA™7 Real-Time PCR System (Life Technologies) and the results assessed with the 12 K Flex QuantStudioTM software (Applied Biosystems). The expression of all genes of interest is presented as fold change relative to sham. Further pathway-focused gene analyses were performed based on the “Enrichr” gene enrichment analysis online tool4_8_-50.

### Ex vivo splenocyte restimulation and multiplex cytokine/chemokine array

Spleens from vaccinated mice for immunological studies are collected at 7 - 10 days post last boost and processed into single cells. Splenocytes are seeded in 24 well plates at 2 x 10^6^ cells per well in 0.5 mL of RPMI media with glutamine, supplemented with 10% fetal bovine serum (FBS) and 100 U/ml penicillin-streptomycin (Invitrogen Life Technologies), and incubated for 72 hours. After 72 hours restimulation, the conditioned media is collected, and the same volume of sterile PBS is added for a two-fold dilution. The conditioned media were profiled using the 32-plex discovery assay (mouse cytokine 32-plex discovery assay; Cat. No: MD32, Eve Technologies Corp., Alberta, Canada). The 32-plex discovery assay consisted of Eotaxin, G-CSF, GM-CSF, IFNγ, IL-1α, IL-1β, IL-2, IL-3, IL-4, IL-5, IL-6, IL-7, IL-9, IL-10, IL-12 (p40), IL-12 (p70), IL-13, IL-15, IL-17, IP-10, KC, LIF, LIX, MCP-1, M-CSF, MIG, MIP-1α, MIP-1β, MIP-2, RANTES, TNFα, and VEGF.

### Challenge studies

Four to five-week-old female BALB/c (H-2d) mice (Charles River Laboratories; Wilmington, MA, USA) were housed at the University of California, Irvine, Vivarium. The University of California, Irvine IACUC approved all animal protocols. The adjuvants CpG-1826 (TriLink, San Diego, CA, USA; 10 μg /mouse/immunization) and Montanide ISA 720 VG (SEPPIC Inc., Fairfield, NJ, USA; 70% of total vaccine volume), FSL-1 were directly mixed with single antigens (MOMP-NLP or MOMP: 10 - 20 μg of each antigen/mouse/ immunization) and antigens combinations (10 μg of each antigen/mouse/immunization). Groups of 5 to 9 mice were immunized twice by the intramuscular (i.m.) route in the quadriceps muscle at a 4-week interval. Immunization controls included EB vaccinations, adjuvant control groups immunized with CpG-1826 and Montanide ISA 720 VG in phosphate buffered saline (PBS). Four weeks after the last immunization mice were challenged intranasally (i.n.) with 10^4^ IFU of *C. muridarum*. All animal experiments were replicated once.

### Statistical analyses

Statistical analyses of secreted cytokines were performed using GraphPad Prism (version 10.1.1) software and comparison between groups was performed using a two-tailed nonparametric Mann–Whitney *t* test. Parametric and non-parametric statistical tests were used as follows for the animal challenge study. The Student’s t-test was employed to evaluate changes in body weight at day 10 p.c., lungs’ weights and amounts of IFN-in lungs supernatants. Repeated measures ANOVA was used to compare changes in mean body weight over the 10 days of observation following the *C. muridarum* i.n. challenge. The Mann–Whitney U-Test was used to compare antibodies titers, levels of IFN- and IL-4 in T-cell supernatants, and the number of *C. muridarum* IFU in the lungs. Values below the limit of detection (BLD) were assigned the value of the BLD, as described by Beal. A *p value* of < 0.05 was considered to be significant. A p value of <0.1 indicates approaching significance.

## Table: Summary of vaccine challenge studies

**Figure 8:**
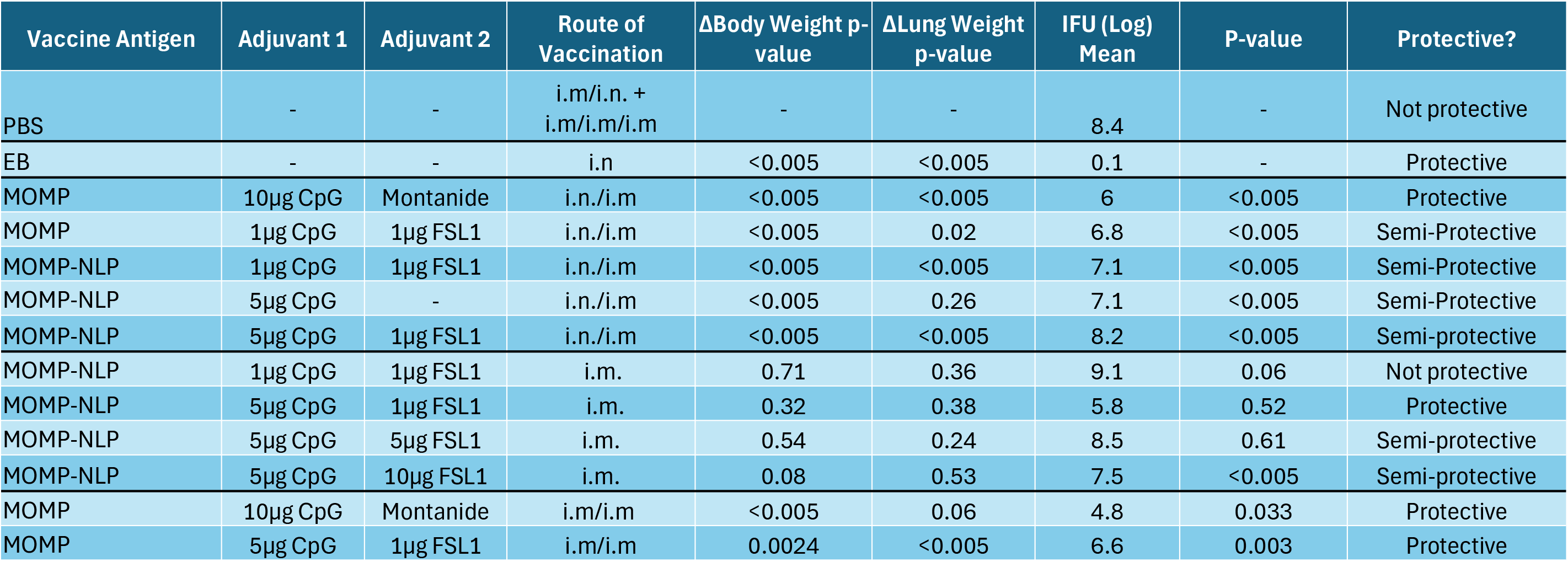
Several *C. muridarum* rMOMP and MOMP-NLP vaccine formulations offer protection against *Chlamydia* infections. Chart summarizing rMOMP and MOMP-NLP formulations tested with various adjuvant combinations and several routes of vaccinations. Protection was determined by 2-fold IFU log mean difference upon comparison to EB values. IFU log means between 1-2 log difference classified as “semi-protective.”Table 1: **Number of GVPs per project** A table of the number of detected GVPs and runs per PRIDE project. For data collected by DDA, the false discovery rate was estimated using the search-then-select procedure and filtered to a 5% peptide-level FDR.

